# Accurate Immune Repertoire Sequencing Reveals Malaria Infection Driven Antibody Lineage Diversification in Young Children

**DOI:** 10.1101/160978

**Authors:** Ben S. Wendel, Chenfeng He, Mingjuan Qu, Di Wu, Stefany M. Hernandez, Ke-Yue Ma, Eugene W. Liu, Jun Xiao, Peter D. Crompton, Susan K. Pierce, Pengyu Ren, Keke Chen, Ning Jiang

**Author notes:** Correspondence should be addressed to: Ning Jiang, Ph.D. Current address: College of Life Sciences, Ludong University, Yantai, Shandong, China, 264025 (Mingjuan Qu). Current address: parasitic Diseases Branch, Division of Parasitic Diseases and Malaria, Center for Global Health (Eugene W. Liu). These authors contributed equally to this work (Ben S. Wendel, Chenfeng He, Mingjuan Qu).

## Abstract

Accurately measuring antibody repertoire sequence composition in a small amount of blood is challenging yet important to the understanding of the repertoire response to infections and vaccinations. Here, we describe an accurate and high-coverage repertoire sequencing method, MIDCIRS, which uses as few as 1,000 naïve B cells. Using it, we studied age-related antibody repertoire development and diversification before and during acute malaria in infants (< 12 months old) and toddlers (12 – 47 months old) with 4-8 ml of blood draws. Unexpectedly, we discovered high levels of somatic hypermutation (SHM) in infants as young as three months old. Antibody clonal lineage analysis revealed that both infants and toddlers increase SHM levels upon infection and memory B cells isolated from pre-malaria samples in malaria-experienced individuals continue to induce SHMs upon malaria rechallenge. These results highlight the vast potential of antibody repertoire diversification in infants and toddlers that has not been realized previously.

## INTRODUCTION

V(D)J recombination creates hundreds of billions of antibodies and T cell receptors that collectively serve as the immune repertoire to protect the host from pathogens. Somatic hypermutation (SHM) further diversifies the antibody repertoire, which makes it impossible to quantify this diversity with nucleotide resolution until the development of high-throughput sequencing-based immune repertoire sequencing (IR-seq)^1,2,3,4^. Although we and others have developed methods to control for artifacts from high amplification bias and sequencing error rates through data analysis^3,5,6,7,8,9^, obtaining accurate sequencing information was only recently made possible by the use of molecular identifiers (MIDs)^10,11,12,13^ MIDs serve as barcodes to track genes of interest through amplification and sequencing. They are short stretches of nucleotide sequence tags composed of randomized nucleotides that are usually tagged to cDNA during reverse transcription to identify sequencing reads that originated from the same mRNA transcript. Despite these advancements, the large amount of input RNA required and low diversity coverage make it challenging to analyze small numbers of cells, such as memory B cells (MemBs) from dissected tissues or blood draws from pediatric subjects, using IR-seq because these samples require many PCR cycles to generate enough material to make sequencing libraries, thus exacerbating PCR bias and errors.

Here we report the development of MID clustering-based IR-seq (MIDCIRS) that further separates different RNA molecules tagged with the same MID. Using naïve B cells (NaiBs), we demonstrate that MIDCIRS has a high coverage of the diversity estimate, or different types of antibody sequences, that is consistent with the input cell number and a large dynamic range of three orders of magnitude compared to other MID based immune repertoire sequencing methods^10,11^. Given the wide use of IR-seq in basic research as well as clinical settings, we believe the method outlined here will serve as an important guideline for future IR-seq experimental designs.

As a proof of principle, we used MIDCIRS to examine the antibody repertoire diversification in infants (<12 months old) and toddlers (12 – 47 months old) from a malaria endemic region in Mali before and during acute *Plasmodium falciparum* infection. Although the antibody repertoire in fetuses^14^, cord blood^15^, young adults^6^, and the elderly^6,16^ has been studied, infants and toddlers are among the most vulnerable age groups to many pathogenic challenges, yet their immune repertoires are not well understood. It is commonly believed that infants have poorer responses to vaccines than toddlers because of their developing immune systems^17^. Thus, understanding how the antibody repertoire develops and diversifies during a natural infection, such as malaria, not only provides valuable insight into B cell ontology in humans, but also provides critical information for vaccine development for these two vulnerable age groups. Using peripheral blood mononuclear cells (PBMCs) from 13 children aged 3 to 47 months old before and during acute malaria, with two of the children followed for a second year and 9 additional pre-malaria subjects, we discovered that infants and toddlers used the same V, D, and J combination frequencies and had similar complementarity determining region 3 (CDR3) length distributions. Although infants had a lower level of average SHM than toddlers, the number of SHMs in reads that mutated in infants was unexpectedly high. Infants showed a similar, if not higher, degree of antigen selection strength, assessed by the likelihood of amino acid-changing SHMs, compared to toddlers. Remarkably, during acute malaria, both infants and toddlers significantly expanded their antibody lineages, and this expansion was coupled with extensive diversification to the same degree as what has been observed in young adults in response to acute malaria^18,19^. Furthermore, informatically reconstructing antibody clonal lineages using sequences from both pre- and acute malaria samples from the same individuals showed that infants were quite capable of introducing SHMs upon a natural infection. This two-timepoint-shared lineage analysis revealed that MemBs isolated from pre-malaria samples in malaria-experienced individuals continued to induce SHMs upon acute malaria rechallenge and most IgM MemBs maintained IgM, while a small fraction switched isotypes. In summary, using an accurate and high-coverage IR-Seq method, we discovered features of the antibody repertoire that were previously unknown in infants and toddlers, shedding light on the development of the immune system and its interactions with pathogens.

## RESULTS

### Overview of the MIDCIRS Method

MIDs have been used to track individual RNA through PCR and sequencing in IR-seq to reduce error rate^10,11,12,13^. They can be designed with sufficient length, and thus diversity, to tag each individual molecule uniquely. However, this requires knowledge of the total number of RNA molecules in the sample, which requires a significant amount of time and effort to measure for each sample before library preparation. Longer MIDs are likely to decrease the reverse transcription efficiency^20,21^. Therefore, we chose 12 random nucleotide MIDs and developed a generalized approach to identify each individual transcript using a sequence-similarity-based clustering method to separate a group of sequencing reads with the same MID into sub-groups. Consensus sequences were then built by taking the average nucleotide at each position within a sub-group, weighted by the quality score. Each consensus sequence represents an RNA molecule, and identical consensus sequences are merged into unique consensus sequences, or unique RNA molecules (Fig. 1a,b and **Supplementary Methods**).

**Figure 1.**
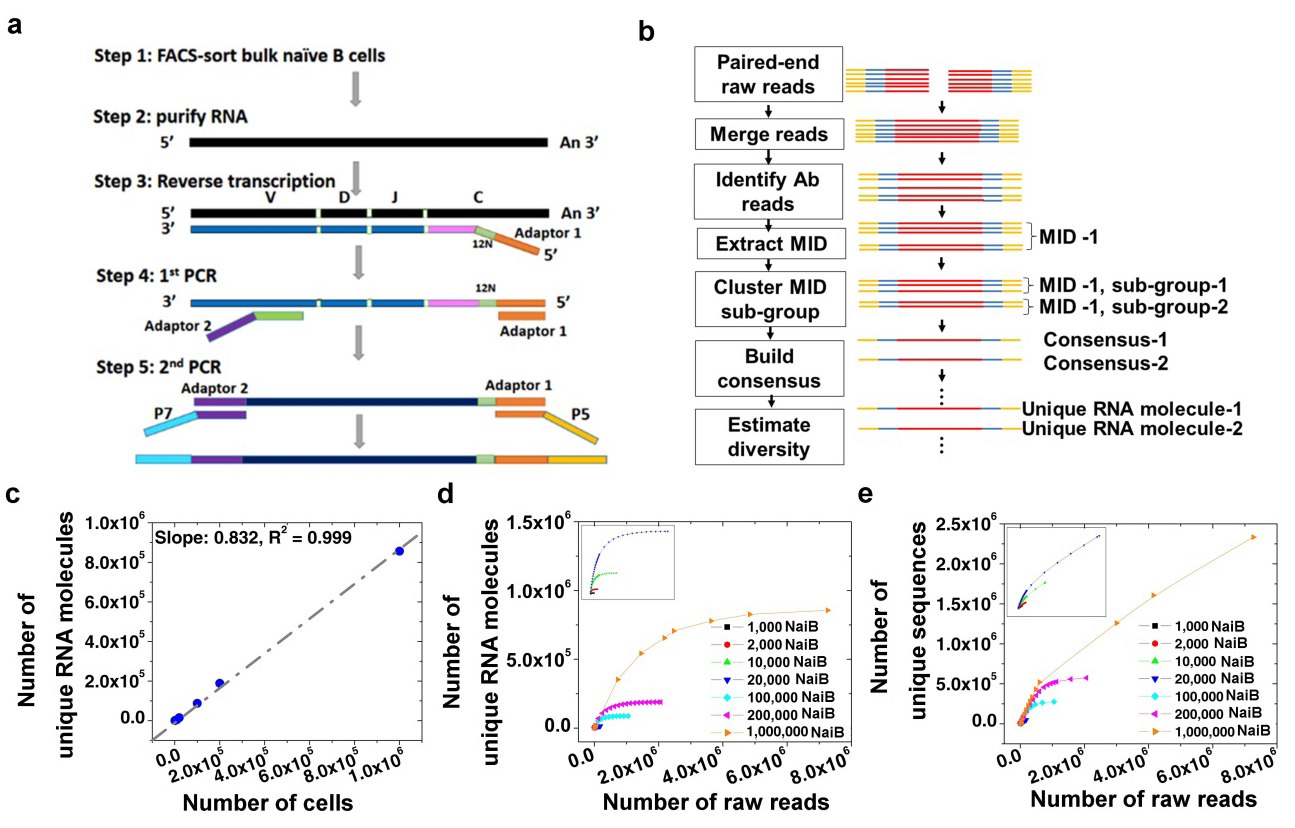
Ultra-accurate high-coverage of antibody repertoire with a large dynamic range of input cells for MIDCIRS. (**a**) Schematic overview of tagging single immunoglobulin transcripts with MIDs. (**b**) Schematic overview of the informatics pipeline of MIDCIRS which includes merging paired-end reads, performing clustering to generate MID sub-groups, and building consensus sequences. (**c**) Correlation between number of cells and number of unique RNA molecules after using MIDCIRS. RNA from as few as 1,000 to as many as 1,000,000 naïve B cells was used as input material in generating the amplicon libraries. Slope indicates the estimated diversity coverage. (**d,e**) Rarefaction analysis of optimum sequencing depth for each sample with (**d**) and without (**e**) using MIDCIRS.

### MIDCIRS yields high accuracy and coverage down to 1000 cells

We used sorted NaiBs with varying numbers (10^3^ to 10^6^) to test the dynamic range of MIDCIRS. The resulting diversity estimates, or different types of antibody sequences, displayed a strong correlation with cell numbers at 83% coverage (Fig. 1c, slope). Previous studies have showed that about 80% of NaiBs express distinct heavy chain genes^22,23^, thus our method achieves a comprehensive diversity coverage that is much higher than other MID-based antibody repertoire sequencing techniques^10,11,12,13^.

We performed rarefaction analysis by subsampling sequencing reads to different amounts and then computing the diversity to test the effect of sequencing depth and error rate on MIDCIRS. On average, the rarefaction curves reached a plateau at a sequencing depth of around three times the cell number using MIDCIRS, suggesting that sequencing more will not discover further diversity (Fig. 1d). In contrast, without using MIDCIRS, the number of unique sequences continued to increase well beyond the number of cells for all samples (Fig. 1e). Optimum sequencing depth is likely to change depending on sample composition (e.g. PBMCs after immunization). Consistent with previous MID-based IR-seq experiments^10^, MIDCIRS reduced the error rate to 1/130^th^ of the Illumina error rate^24^, providing the accuracy necessary to distinguish genuine SHMs (1 in 1,000 nucleotides^25^) from PCR and sequencing errors (1 in 200 nucleotides^24^) (**Supplementary Fig. 2**).

### Infants and toddlers have similar VDJ usage and CDR3 lengths

Equipped with this ultra-accurate and high-coverage antibody repertoire sequencing tool, we applied it to study the antibody repertoire of infants and toddlers residing in a malaria endemic region of Mali. From an ongoing malaria cohort study^26^, we obtained paired PBMC samples collected before and during acute febrile malaria from 13 children aged 3 to 47 months old (**Supplementary Fig. 3** and **Supplementary Table 3**). Two of the children were followed for an additional year, giving 15 total paired PBMC samples. An average of 3.8 million PBMCs per sample were directly lysed for RNA purification. All PBMCs were subjected to MIDCIRS analysis. An average of 3.75 million sequencing reads were obtained for each PBMC sample (**Supplementary Table 4**).

For all PBMC samples, sequencing approximately the same number of reads as the cell numbers saturated the rarefaction curve (**Supplementary Fig. 4**). VDJ gene usage was highly correlated for IgM between infants and toddlers regardless of weighting the correlation coefficient by the number of sequencing reads or clonal lineages (**Supplementary Fig. 6**), demonstrating that the same mechanism of VDJ recombination is used to generate the primary antibody repertoire in infants and toddlers. Increased correlation was also seen for IgG and IgA when weighting on the number of clonal lineages in each VDJ class; however, the correlation coefficient was lower when weighting on the number of reads in each VDJ class (**Supplementary Fig. 6**). The diagonal lines in each panel indicate same sample self-correlation, and the two shorter off-diagonal lines indicate correlations from two timepoints of the same subject. These data recapitulate previous observations from our study in zebrafish that clonal expansion-induced differences on the number of reads in each VDJ class can confound the highly similar VDJ usage during B cell ontology^5^. In addition, our results of similar CDR3 length distribution in both infants and toddlers across the three isotypes and both timepoints (**Supplementary Fig. 7**) are consistent with recent studies of PBMCs of 9 month olds infants^14,15^ and adult PBMCs^27,28^ and confirms the previous results that an adult-like distribution of CDR3 length is achieved around two months of age^29^.

### Both infants and toddlers have unexpectedly high SHMs

SHM is an important characteristic of antibody repertoire secondary diversification due to antigen stimulation^30^. Although it has been demonstrated before that infants have fewer mutations in their antibody sequences compared to toddlers and adults, the limited number of sequences for only a few V genes did not provide convincing evidence of the levels of SHM in infants^31^. A recent study using the first generation of IR-seq showed that they average at least 6 SHMs in IgM of an average length of 500 nucleotides increased in two 9 months old infants^14^. These numbers are equivalent to, if not higher than, what have been reported for SHM rate observed in IgMs in healthy adults day 7 post influenza vaccination^10^ and was much higher compared to another study using a few V genes and limited antibody sequences^32^. Due to inherent errors associated with the first generation of IR-seq as discussed above, it is possible that PCR and sequencing errors played a role here^14^. In addition, it remains unclear if infants (< 12 months old) are able to generate a significant number of mutations in response to infection, which would demonstrate their capacity to diversify the antibody repertoire^33^.

We discovered that infants (< 12 months old) and toddlers (12 – 47 months old) had an unexpectedly high level of SHMs in all three major isotypes of antibodies (Fig. 2a) when plotting the histogram of all unique RNA sequences according to the number of SHMs. While mutation distribution remained in the low end of the spectrum for IgM in infants and toddlers, mutations were significantly higher in IgG and IgA for all age groups, with mutation threshold of top 10% unique RNA molecules increasing to around 10 in infants (**Fig.2a**, Inf, right of the blue long vertical lines) and almost 20 in toddlers (Fig. 2a, Tod, right of the blue long vertical lines). Our analysis also showed a previously unappreciated high level of mutations in IgA^34^. To minimize any possible inflation of SHMs, we excluded all sequences that were mapped to novel alleles, which were identified by both TIgGER^35^ and inspecting the IgM sequences in PBMC samples (**Supplemental Methods**). These putative novel alleles account for 8% of all unique sequences on average (**Supplementary Table 5**). We also analyzed SHM in sorted NaiBs as a control, which showed extremely low SHM load, 0.55 mutations on average, as expected (**Supplementary Table 6**). Upon an acute malaria infection, the SHM histogram shifted rightward for almost all isotypes in almost all individuals (Fig. 2a, the right shift of pink long vertical line compared to blue long vertical line), including infants. These results demonstrated unexpected high level of SHM that exceed what have been documented previously^14^.

**Figure 2.**
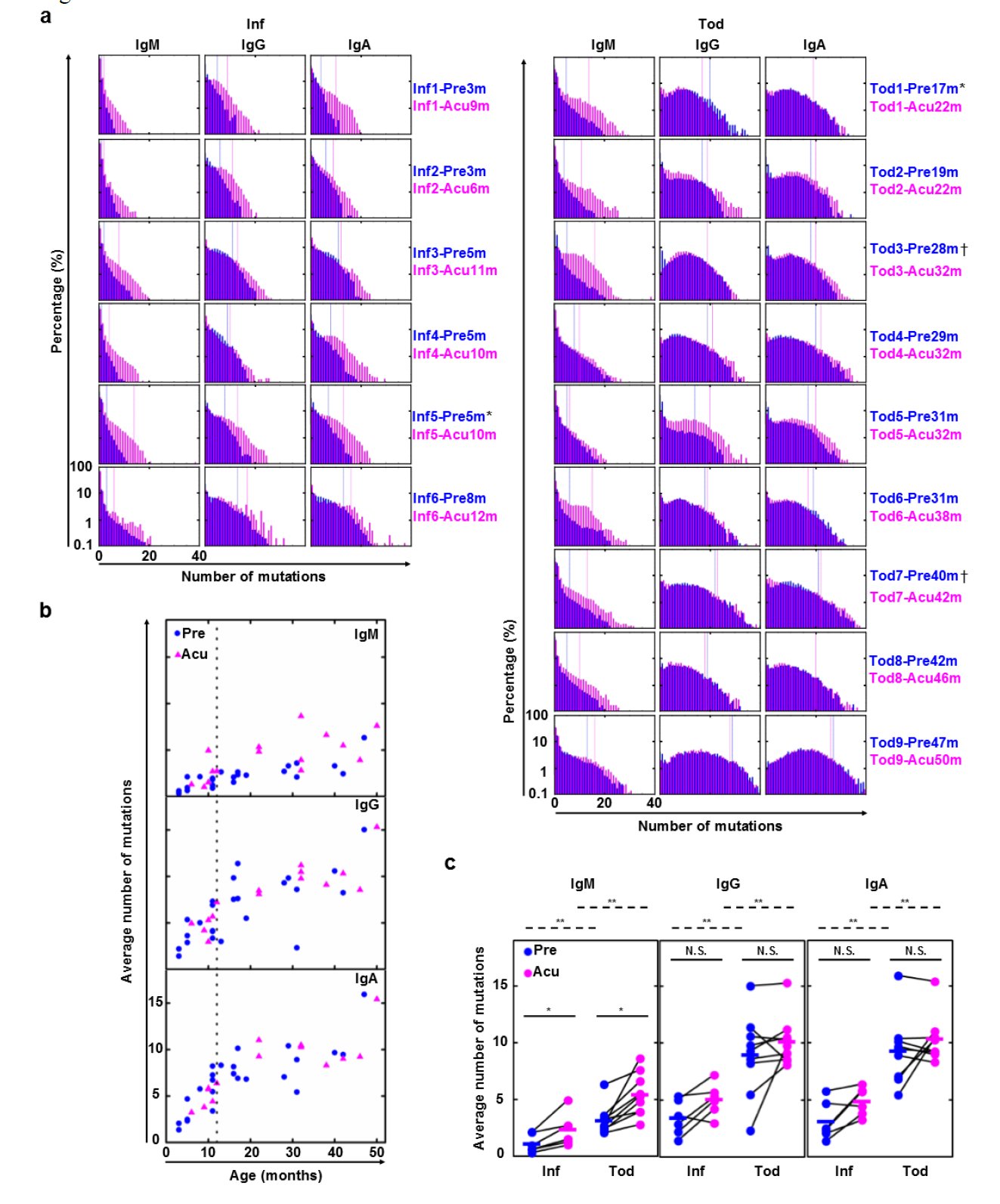
Infants and toddlers are separated into two stages based on SHM load. (**a**) Distribution of SHM number for infants (N=6) and toddlers (N=9), from whom we had paired pre- (blue) and acute (pink) malaria samples, weighted by unique RNA molecules. Blue and pink long vertical lines represent the number of mutations above which 10% of sequences fall for the respective samples. ^*^ and † demarcate samples derived from the same individuals followed for 2 malaria seasons. (**b**) Age-related average number of mutations in pre- (blue circle, N=24, N_Infant_=11, N_Toddler_=13) and acute malaria (pink triangle, N=15, N_Infant_=6, N_Toddler_=9) samples, weighted by RNA molecules. Dashed line indicates the age boundary for infants (<12 months old) and toddlers (12 – 47 months old). (**c**) Comparison of average number of mutations for paired infants and toddlers. Pre- (blue) and acute (pink) malaria samples separated by isotype; lines connect paired samples (N_Infant,paired_=6, N_Toddler,paired_=9). Bars indicate means. ^*^*P <* 0.05, ^**^*P* < 0.01, N.S. indicates no significant difference by two-tailed Mann-Whitney U test (between age groups, dashed lines) or two-tailed Wilcoxon Signed-Rank test (between paired timepoints, solid lines). Differences in variance were not significant by squared ranks test.

### SHM load is distinct between infants and toddlers

The differences of SHM distributions between infants and toddlers, decreasing mutation histogram from 0 for infants in all three isotypes while peaking at 10 for toddlers in IgG and IgA (Fig. 2a), suggest that the total SHM load might reflect the history of interactions between their antibody repertoires and the environment, including malaria exposure. Since the malaria season is synchronized with the 6-month rainy season (**Supplementary Fig. 3)** and > 90% of subjects in this cohort are infected with *P. falciparum* during yearly malaria season^26^, we hypothesized that the SHM load would increase with increasing age. However, we found that the SHM load rapidly increased with age in infancy and then appeared to plateau around 12 months of age in an initial smaller set of subjects with paired pre- and acute malaria paired PBMC samples (**Supplementary Fig. 8**). We then added 9 pre-malaria samples around the infant and toddler transition (5 of 11 months old and 4 of 13 to 17 months old). This two-staged trend of SHM load remained for all three isotypes (Fig. 2b), with samples around the transition having the largest variation. Detailed comparisons showed that, consistent with the two-stage trend, toddlers had a higher SHM load compared to infants for all three isotypes at both pre- and acute malaria (Fig. 2c, comparison between age groups). Although there was significant increase on SHM load upon an acute malaria infection in IgM for both infants and toddler, bulk PBMC analysis did not show a significant increase in IgG or IgA, possibly because of an already elevated SHM base level. This, along with the two-stage trend we saw (Fig. 2b), suggests that 12 months is an important developmental threshold for secondary antibody repertoire diversification: before this threshold, the global repertoire is quite naïve but can quickly diversify upon a natural infection.

### Higher MemB percent in toddlers results in a higher SHM load

This unexpected developmental threshold of secondary antibody repertoire diversification prompted us to focus on B cell subset composition changes and ask whether they correlate with this two-staged SHM load. Flow cytometry analysis revealed that NaiBs decreased from about 95% in 3-month-old infants to about 80% in toddlers (Fig. 3a). Conversely, MemBs increased from about 4% in 3-month-old infants to about 15% in toddlers (Fig. 3f). As is suggested from the two-stage SHM load analysis, 12 months appears to divide the samples into two age groups, with a large variation at the infant to toddler transition and in the toddler group. Percentages of NaiB and MemB in the infant group were significantly different than those in the toddler group (Fig. 3b,g). Plasmablast percentages fluctuated in a much smaller range (**Supplementary Fig. 10**). With similar two-staged trend observed for B cell subset percentages, we hypothesized that the B cell subset percentage would correlate with SHM load. Indeed, further analysis showed that the decrease in NaiB percentage and the increase in MemB percentage correlated well with SHM load across IgM, IgG, and IgA isotypes (**Fig. 3c-e** and **h-j**), which supports our initial hypothesis that 12 month separates infants from toddlers in both SHM load and B cell composition changes. These data suggest that MemBs contribute significantly to the developing antibody repertoire, and their composition is essential in secondary antibody repertoire diversification.

**Figure 3.**
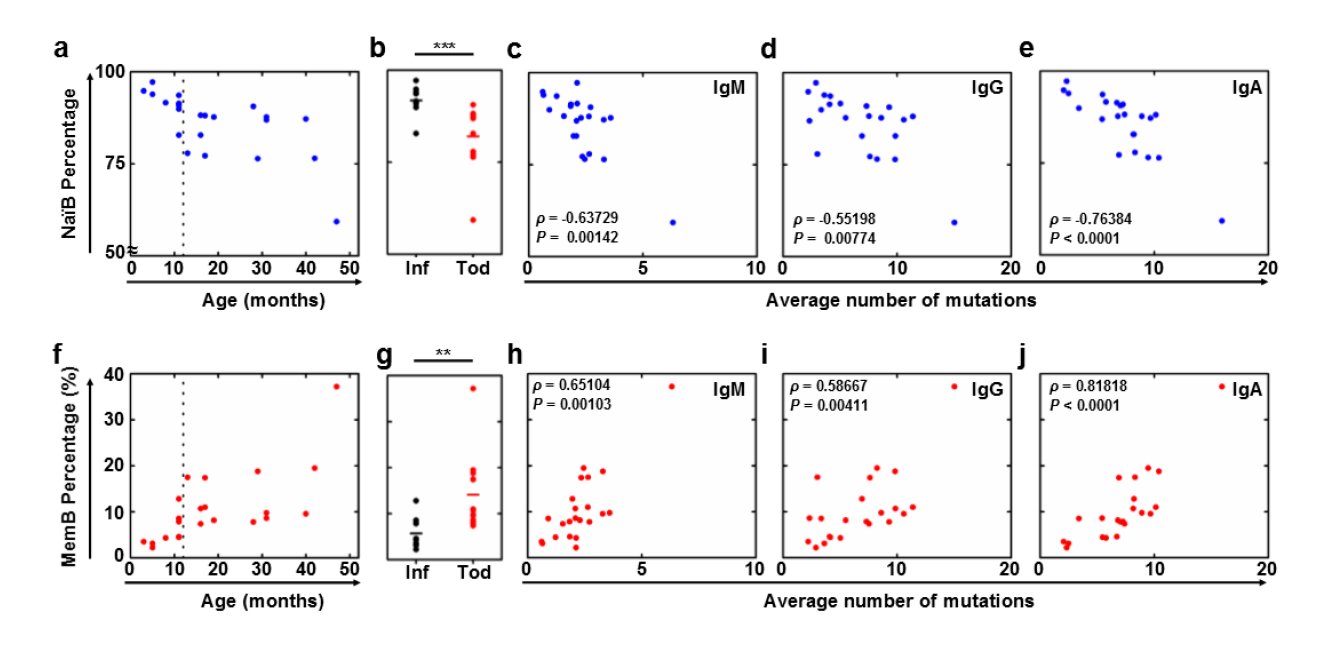
Decrease of naïve B cell and increase of memory B cell percentages show a two-stage trend and correlate with SHM load. (**a**) NaïB percentages of total B cells from the pre-malaria samples (N=22) vary with age. Dashed vertical line depicts the cutoff between infants and toddlers. (**b**) NaïB percentages of total B cells compared between infants (black, N=9) and toddlers (red, N=13). (**c-e**) NaïB percentages correlate with average number of mutations (SHM load) in IgM (**c**), IgG (**d**), and IgA (**e**) sequences from bulk PBMCs in pre-malaria samples (N=22). (**f**) MemB percentages of total B cells from the pre-malaria samples (N=22) vary with age. Dashed vertical line depicts the cutoff between infants and toddlers. (**g**) MemB percentages of total B cells compared between infants (black, N=9) and toddlers (red, N=13). (**h-j**) MemB percentages correlate with average number of mutations (SHM load) in IgM (**h**), IgG (**i**), and IgA (**j**) sequences from bulk PBMCs in pre-malaria samples (N=22). (**b** and **g**) Bars indicate means; ^**^*P* < 0.01, ^***^*P* < 0.001, two-tailed Mann-Whitney U test. (**c** to **e** and **h-j**) *ρ* and *P* values determined by Spearman’s rank correlation listed in each panel.

### SHMs are similarly selected in infants and toddlers

One of the key features of antibody affinity maturation is antigen selection pressure imposed on an antibody, which is reflected in the enrichment of replacement mutations^36^ in the CDRs, the parts of the antibody that interact with antigens, and the depletion of replacement mutations in the framework regions (FWRs), the parts of the antibody responsible for proper folding. The unexpectedly high level of SHMs observed in infants prompted us to ask whether those SHMs that occurred in infants have characteristics of antigen selection, as seen in older children and adults. As previous studies have shown that infants have limited CD4 T cell responses and neonatal mice exhibit poor germinal center formation^17^, we hypothesized that infant antibody sequences would display weaker signs of antigen selection. To this end, we used a recently published tool, BASELINe^37^, to compare the selection strength. BASELINe quantifies the likelihood that the observed frequency of replacement mutations differs from the expected frequency under no selection; a higher frequency implies positive selection and a lower frequency implies negative selection, and the degree of divergence from no selection relates to the selection strength. Surprisingly, despite infants harboring fewer overall mutations, these mutations were positively selected in the CDRs and negatively selected in the FWRs in both IgG and IgA (Fig. 4b,c,e,f). Contrary to the hypothesis that infants would have a lower selection strength compared to toddlers, for both IgG and IgA, infants actually had a higher selection strength at both pre- and acute malaria compared to toddlers (Fig. 4). The lower selection strength in infant IgM at pre-malaria was significantly increased at acute malaria (Fig. 4a,d, CDR black curves between two timepoints, *P* < 0.0001, as previously described^37^), suggesting that the significant increase in SHM was antigen-driven and selected upon. In order to compare with a large amount of historical adult data, we calculated replacement to silent mutation ratio (R/S ratio), which were about 2-3:1 in FWRs and 5:1 in CDRs for both infants and toddlers (**Supplementary Table 7**). These results were similar to adults^36,37,38,39,,40^ and much higher than what has been reported for children previously using a very limited number of sequences^41^. We also noticed that R/S ratio in the FWRs of IgM was much higher in infants, contrary to the BASELINe results, which highlights the importance of incorporating the expected replacement frequency when considering selection pressure. These results suggest that as an end result of interactions between antigen selection and SHM, the degree of antibody amino acid changes is comparable in infants, toddlers, and adults. It also suggests that cellular and molecular machineries for antigen selection are already in place in infants.

**Figure 4.**
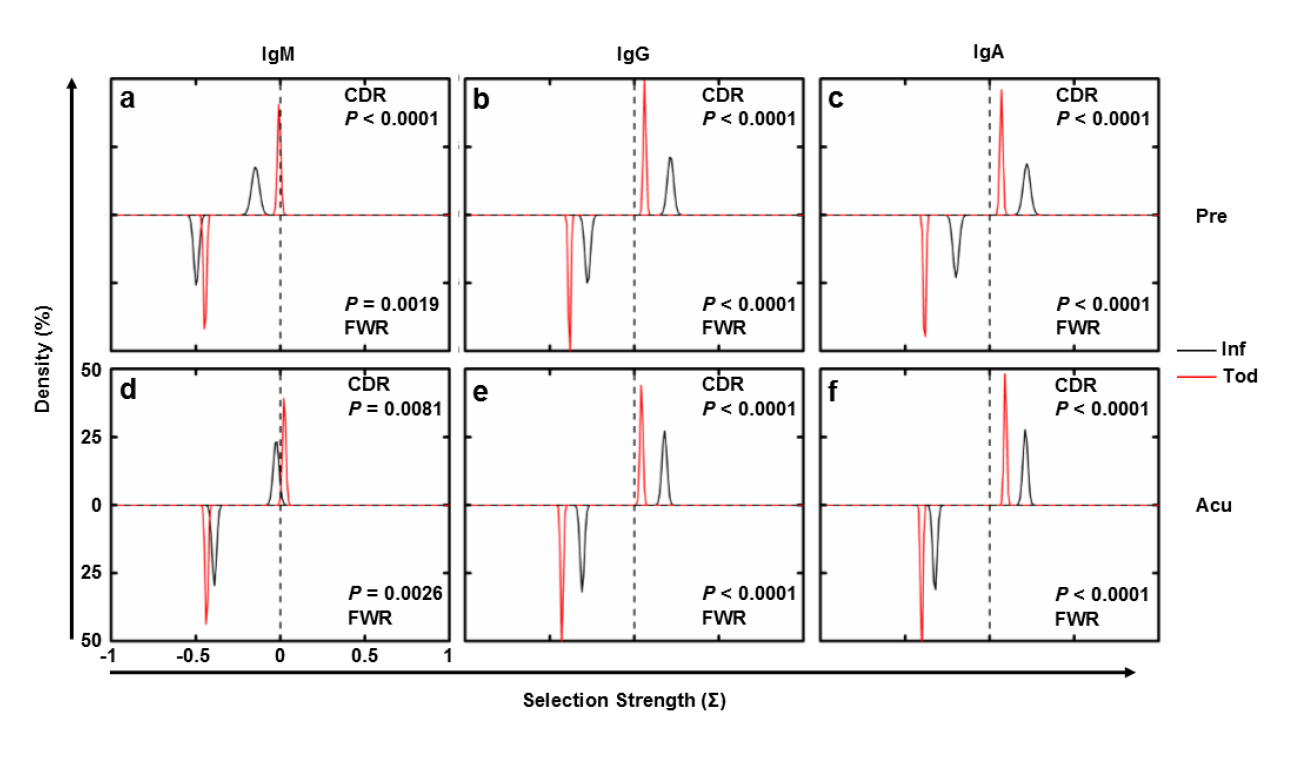
Antigen selection strength comparisons between infants and toddlers. Selection strength distributions, as determined by BASELINe^37^, were compared between infants (black) and toddlers (red) for PBMCs from pre- (**a-c**) (N_infant_=6, N_toddler_=9) and acute (**d-f**) (N_infant_=6, N_toddler_=9) malaria timepoints, separated by isotype: (**a,d**) IgM, (**b,e**) IgG, and (**c,f**) IgA. Selection strength on CDR (CDR1 and 2, top half of each panel) and FWR (FWR2 and 3, bottom half of each panel) for unique RNA molecules was calculated. CDR3 and FWR4 were omitted due to the difficulty in determining the germline sequence. FWR1 for all sequences was also omitted because it was not covered entirely by some of the primers. *P* value calculated as previously described^37^.

### Clonal lineages diversify upon an acute febrile malaria

The exhaustive sequencing data obtained by MIDCIRS offered the possibility to reconstruct the clonal lineages that trace B cell development. Clonal lineages contain different species of unique antibody sequences that could be progenies derived from the same ancestral B cell. B cell clonal lineage analysis has been used to track affinity maturation and sequence evolution of HIV broadly neutralizing antibodies^42,43^. Using a clustering method with a pre-determined threshold (90% similarity on nucleotide sequence at CDR3), we previously demonstrated that B cell clonal lineages could be informatically defined and contain pathogen-specific antibody sequences^6^. In addition, the clonal lineage analysis also highlighted the lack of antibody diversification in the elderly after influenza vaccination^6^. Using the same approach and a similar threshold^6,44^, we aimed to answer these questions – whether infants and toddlers are able to diversify antibody clonal lineages in response to infection and, if so, whether they have a similar ability to do so, which were previously impossible to answer due to technical limitations. To do this, we first visualized structures of informatically defined clonal lineages for the entire antibody repertoire (**Supplementary Fig. 11**). Each oval lineage map represents an individual PBMC sample at one timepoint. Densely packed individual lineages are not easily identified visually in **Supplementary Fig. 11**; however, dark areas indicate that clonal lineages are already complex in this cohort of infants as young as 3 months old and can be further diversified upon a febrile malaria.

The densely packed lineages could result from large lineage sizes (one unique RNA molecule with many copies), large lineage diversities (many unique RNA molecules), or a combination of the two. To closely examine the possible differences in the degree of this intra-clonal lineage expansion and diversification between infants and toddlers, especially upon acute febrile malaria, we projected the global lineage structure (**Supplementary Fig. 11**) onto diversity and size of lineage axes (Fig. 5a). Each circle represents an individual lineage, with the area of the circle proportional to the SHM load (average mutations of the lineage). This analysis effectively captures five parameters that quantify lineage complexity in a sample: number of total clonal lineages (number of circles), diversity of each lineage (x-axis position, number of unique RNA molecules in a lineage), size of each lineage (y-axis position, number of total RNA molecules in a lineage), SHM load of each lineage (area of circle, key is located in between the infant and toddler panels in Fig. 5a), and the extent of clonal expansion of each lineage (distance from y=x parity line; no clonally expanded RNA molecules within a lineage if it is on parity line or pure clonal expanded RNA molecules if it is in the top left quadrant of each panel).

**Figure 5.**
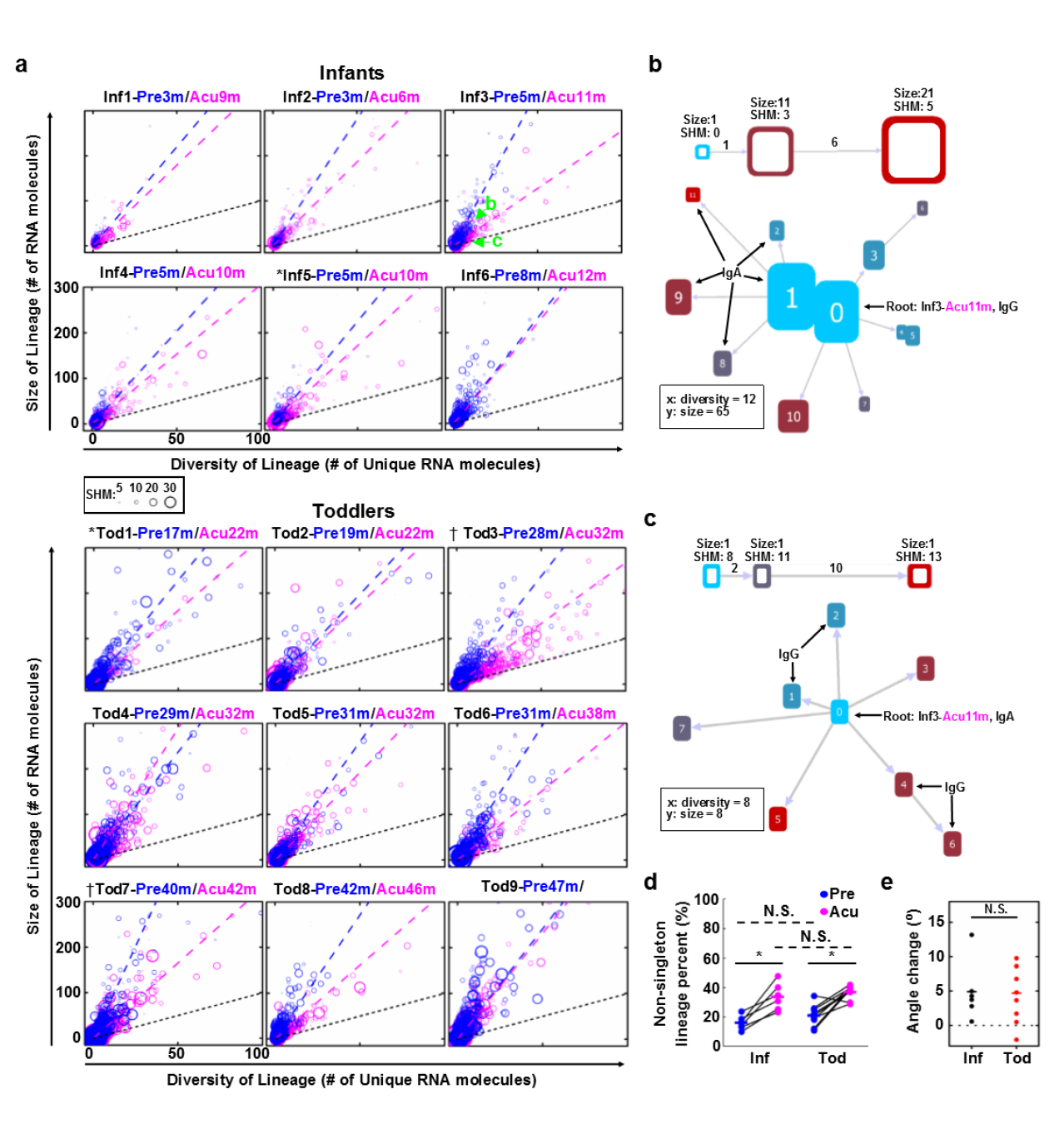
B cell lineage complexity change under malaria stimulation. (**a**) Diversity and size of B cell lineages for infants (N=6) and toddlers (N=9) from whom paired PBMC samples at pre- (blue) and acute (pink) malaria were obtained. Each circle represents an individual lineage. The area of each circle is proportional to the SHM load. Labeled green arrows indicate representative lineages whose intra-lineage structures were shown in detail in (**b**) and (**c**). Each circle’s x and y coordinates were determined by its diversity (the number of unique RNA molecules in a lineage) and size (the number of total RNA molecules in a lineage), respectively. Blue and pink dashed lines represent the linear fit for pre- and acute malaria lineages, respectively. Black dashed lines indicate y=x parity, such that lineages lying on the parity line are comprised entirely of unique RNA molecules with minimum clonal expansion, such as lineage in (**c**). On the other hand, lineages comprised of clonally expanded RNA molecules are close to the y axis, such as lineage (**b**). (**b,c**) Each node is a unique RNA molecule species. The height of the node corresponds to the number of RNA molecules of the same species, the color corresponds to number of nucleotide mutations, and the distance between nodes is proportional to the Levenshtein distance between the node sequences, as indicated in the legend above each lineage. All unlabeled nodes share the isotype with the root. (**d**) The non-singleton lineage percent (lineages comprised of at least 2 RNA molecules) between infants and toddlers at pre- (blue) and acute (pink) malaria. ^*^*P* < 0.05 by two-tailed Wilcoxon Signed-Rank test (between timepoints, solid lines); N.S. indicates no significant difference by two-tailed Mann-Whitney U test (between age groups, dashed lines). (**e**) The difference of linear regression slopes (angles), or degree of diversity change, between pre- and acute malaria for infants (black) and toddlers (red). N.S. indicates no significant difference by two-tailed Mann-Whitney U test. Bars indicate means. Differences in variance were not significant by squared ranks test.

Fig. 5b,c are two example lineages selected to display the full lineage structures to demonstrate a lineage with diversification and clonal expansion (Fig. 5b refers to green letter “b” indicated in Fig. 5a, Inf3) and another one with diversification but without clonal expansion (Fig. 5c refers to green letter “c” indicated in Fig. 5a, Inf3). Both were represented by a single circle in Fig. 5a, but their locations in Fig. 5a depend on their numbers of RNA molecules (y-axis) and numbers of unique RNA molecules (x-axis). Lineage “c” (c in Fig. 5a, Inf3, zoomed in view in Fig. 5c) that lies away from the origin and near the black y=x parity line consists of 8 unique sequences, each represented by only one RNA molecule, indicating extensive lineage diversification but no clonal expansion. Lineage “b” (b in Fig. 5a, Inf3, zoomed in view in Fig. 5b) that lies far from the parity line is dominated by two unique RNA molecules each with about 20 copies (Fig. 5b, height of nodes), indicating extensive clonal expansion of particular sequences in addition to diversification. Changing lineage forming threshold from 90% to 95% did not change the overall structure of the lineages (**Supplementary Fig. 12**).

This five-dimension lineage analysis revealed that infants as young as 3 months old can generate extensive lineage structures, with many lineages containing more than 20 different types of antibody sequences and 50 RNA molecules (Fig. 5a). Toddlers had many more lineages with higher levels in both size and diversity. However, in both infants and toddlers, the majority of these clonal lineages were singleton lineages consisting of only one RNA molecule (Fig. 5d), consistent with the flow cytometry analysis that the bulk of the B cell repertoire was naïve in these young children (Fig. 3). Upon acute malaria infection, the fraction of non-singleton lineages increased in both infants and toddlers **(Fig. 5d)**.

In order to tease out whether these non-singleton lineages diversify or clonally expand upon acute infection, we fitted linear regressions to the lineage diversity-size plot. An immune response against an infection can have a two-fold effect on the lineage landscape: antigen stimulation can cause clonal expansion, which would shift the lineage up on the y-axis, and SHM and affinity maturation, which would shift the lineage to the right on the x-axis. This balance between clonal expansion and diversification is depicted by the slope of the linear regression (Fig. 5a, dashed blue lines for pre-malaria samples and dashed pink lines for acute malaria samples). We hypothesized that the lower absolute SHM load of infants would imply a defect in the ability to diversify clonal lineages in response to infection, leading the slope change from pre- to acute malaria to be low (a small angle between blue and pink dashed lines) or even negative (pink dashed line is closer to y-axis compared to blue dotted line). Surprisingly, we found that infants diversify their clonal lineages in a similar manner as toddlers in response to acute malaria (Fig. 5e). As singleton lineages do not bear any weight on the linear regression, our analysis showed that the increased fraction of non-singleton lineages upon malaria infection were similarly diversified between infants and toddlers, which was also similar to what was observed in a young adult at pre- and acute malaria (**Supplementary Fig. 14**). However, this sharply contrasts with what we had previously observed in the elderly following influenza vaccination, where clonal expansion dominated^6^.Among clonally expanding and diversifying B cell clones during an infection, only a subset of the cells comprising the clonal burst remain once the infection has been cleared. Thus, the characteristic change in the lineage size/diversity linear regression slope upon infection is expected to subside as time passes since the acute infection. Indeed, comparing the pre-malaria lineage size/diversity linear regression slopes revealed no difference between infants (who have not experienced malaria before) and toddlers (who have experienced malarias in previous years) (**Supplementary Fig. 13**). These results highlight the unexpected capability of young children’s antibody repertoire in response to a natural infection.

### SHM load increases upon an acute febrile malaria infection

The plateau we observed on SHM load in toddlers at both pre- and acute malaria (Fig. 2b) and the lack of a SHM difference in IgG and IgA between pre- and acute malaria (Fig. 2c) seems to suggest that the experienced part of the repertoire did not respond to malaria infection by inducing SHM. However, it could be that only a portion of the bulk antibody repertoire responded to the infection and there was already a high level of baseline SHMs as revealed by the histogram analysis (Fig. 2a). Since we saw the lineage diversification upon malaria infection in Fig. 5, we hypothesized that examining the SHMs from sequences in the two-timepoint-shared lineages (lineages containing both pre- and acute malaria sequences) would enable us to quantify the infection induced SHM increase from the highly mutated background. To test this, we pooled all sequences from both timepoints, including sorted MemBs at pre-malaria, and generated lineages again using 90% similarity threshold at CDR3^6,44^. We were able to find two-timepoint-shared lineages in all subjects analyzed (**Supplementary Table 8**). Consistent with the other observation that toddlers already have diversified antibody repertoire compared to infants, there are more shared lineages in toddlers than in infants (**Supplementary Table 8**). We tallied SHMs for sequences from pre- and acute malaria in the two-timepoint-shared lineages separately. Consistent with our hypothesis, both infants and toddlers significantly increased SHM upon infection (Fig. 6a). Indeed, toddlers had a higher pre-malaria SHM level compared to infants (Fig. 6a, blue symbols). To our surprise, infants were able to induce more SHMs compared to toddlers (Fig. 6b). These data suggested that indeed both infants and toddlers induce SHMs upon malaria infection.

**Figure 6.**
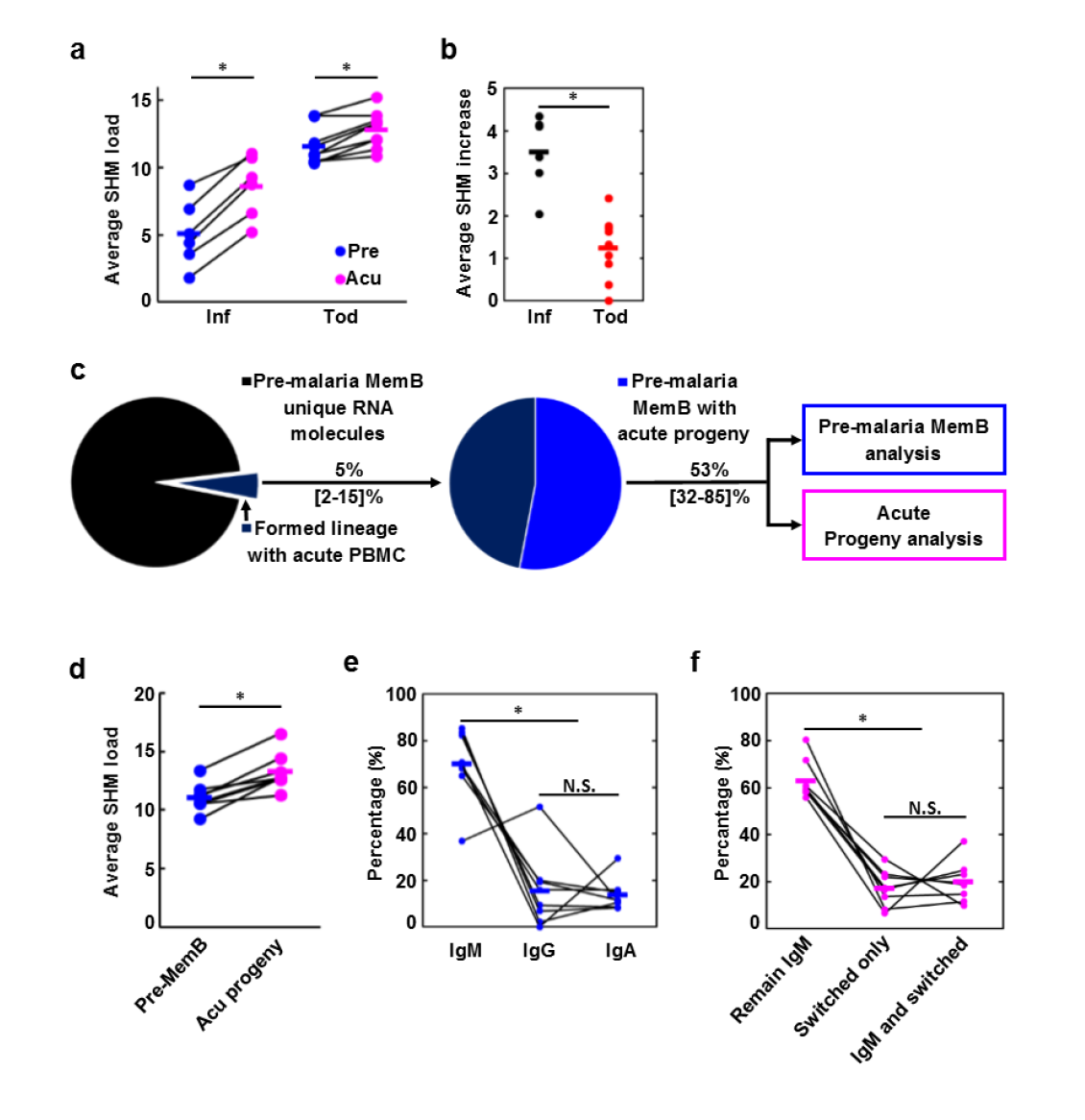
Two-timepoint-shared lineage analysis reveals SHM increment during acute malaria infection. (**a**) Average SHM for sequences from pre- (blue) and acute (pink) malaria timepoints within lineages containing sequences from both timepoints for infants (N=6) and toddlers (N=9). (**b**) Average SHM increase upon acute malaria infection for infants (black) and toddlers (red) from (**a**). (**c**) Flow diagram for two-timepoint-shared lineage containing pre-malaria MemB identification and acute progeny analysis. Percentages represent the average percent of unique sequences classified by the indicated slice, range in brackets. (**d**) Average SHM load for pre-malaria MemBs with acute progeny (blue) and their acute progenies (pink) for malaria-experienced toddlers with FACS sorted pre-malaria MemBs (N=8). (**e**) Isotype distribution of pre-malaria MemBs with acute progeny. (**f**) Isotype fate of acute progenies stemming from IgM pre-malaria MemBs. Lines connect the same subjects. Bars indicate means. (**a**, **d-f**) ^*^*P* < 0.05, N.S. indicates not significant by two-tailed Wilcoxon Signed-Rank test. (**b**) ^*^*P <* 0.05 by two-tailed Mann-Whitney U test.

### Pre-malaria MemBs further diversify upon malaria rechallenge

The importance of IgM-expressing MemBs has been reported in mice in several recent studies^45,46,47,48^, including a mouse model of malaria infection^49^. However, fewer studies have examined these cells in humans^50^, and their composition and role in repertoire diversification upon rechallenge remains elusive. Many suspect that they may retain the capacity to introduce further mutations and class switch^45,46,49,51^. However, sequence-based clonal lineage evidence is lacking. The paired samples before and during acute malaria from malaria experienced toddlers provided an opportunity to investigate the role of MemBs in repertoire diversification upon rechallenge in children.

Here, we focused on two-timepoint-shared lineages that harbored sequences from pre-malaria MemBs. Given the significant increase of SHM we identified at acute malaria sequences over pre-malaria sequences in two-timepoint-shared lineages (Fig. 6a), we reasoned that the high repertoire coverage of MIDCIRS should enable us to identify a large number of two-timepoint-shared lineages that contain these MemBs, and these MemBs should have mutated progenies at the acute malaria timepoint. To ensure that we identify sequence progenies of these pre-malaria MemBs, we employed an antibody lineage structure construction algorithm, COLT, that we recently developed^52^. COLT considers isotype, sampling time, and SHM pattern when constructing an antibody lineage, which allows us to trace, at the sequence level, the acute progeny of these MemBs. As illustrated by **Supplementary Fig. 15**, this COLT-generated lineage tree depicts a pre-malaria MemB sequence serving as a parent node to sequences derived from the acute malaria timepoint. This analysis is much more stringent in identifying sequence progenies than simply judging if a pre-malaria MemB sequence is grouped with acute malaria PBMC sequences.

On average, 5% of unique sequences from 10,000 sorted MemBs formed lineages with acute malaria PBMC sequences (Fig. 6c, dark blue slice of the first pie). We performed COLT^52^ analysis on these pre-malaria MemB-containing lineages and found that 53% contained traceable progeny sequences from the acute malaria PBMCs (Fig 6c, lighter blue slice of the second pie). Overall, there is a significant increase on SHM in these acute malaria progenies compared to their ancestor pre-malaria MemBs (Fig. 6d). Consistent with previous studies^45,46,51^, these progeny-bearing pre-malaria MemBs expressed all three isotypes, with IgM being the dominant species (Fig. 6e). We also investigated their isotype switching capacity and found that about 60% of the IgM pre-malaria MemBs maintained IgM as progenies; however, about 20% only had isotype-switched progenies detected while the remaining 20% had both IgM and isotype switched progenies (Fig. 6f). These pre-malaria IgM MemBs largely retain IgM expression while further introducing SHM upon rechallenge. Thus, these analyses showed multi-facet diversification potential of young children’s MemB in a natural infection rechallenge.

## DISCUSSION

About 13,000 children under one year old die every day worldwide^53^, and most of these deaths were caused by infection^17^. It has long been recognized that children’s immune systems are immature at birth and require time to develop to provide protection against pathogens or respond to vaccines. However, few studies have focused on children’s antibody repertoire development, diversification, and response to infection. Knowledge in this area holds great interest to vaccine development and vaccination strategy design. This is especially urgent for malaria, as it still kills about half a million children each year^54^, and the most advanced malaria vaccine confers only partial, short-lived protection in African children^55^.

However, studying the antibody repertoire in young children is challenging in several regards: first, lack of analytical tools to exhaustively study the antibody repertoire from small volumes of blood, second, lack of informatic analysis tools to turn high-throughput data into knowledge, and third, the rarity of a large set of samples from young children obtained before and at the time of a natural infection. To address these challenges, we developed a highly accurate and high-coverage immune repertoire sequencing tool, MIDCIRS, to analyze antibody repertoire development, diversification, and capacity to respond to a natural infection in children who were experiencing acute febrile malaria.

Previous studies showed that there was evidence of SHM and antigen selection in infants 8 months of age or older by examining a few V gene alleles^31^. However, it is not clear how widespread SHMs are in infant antibody repertoires and to what degree SHMs can be introduced in response to an infection. By using a comprehensive and unbiased analysis, we showed that infants as young as 3 months old can have 10% of sequences with 5 or more mutations, and they can further introduce mutations upon an acute febrile malaria to well over 20 SHM per 270nt heavy chain V region. Compared to toddlers, there is a separation on SHM load around 12 months: this number gradually increases before 12 months and stays at a plateau after that regardless of repeated malaria incidents. Consistent with this trend is the similar pattern observed in the increase in the percentage of MemB and corresponding decrease in percentage of NaiB with age: both plateaued after 12 months of age. Accordantly, SHM load in IgM, IgG, and IgA correlated with the percentage of NaiB and MemB. Surprisingly, regardless of the lower mutation load in infants, their mutations were similarly, if not more strongly, selected as those of toddlers, suggesting that the molecular machineries and other cellular components involved in antibody selection are already developed in infants. In future analysis, it will be of interest to tease out the mechanistic contributions to a two-stage increase of average mutation number, in particular the role of T cell help and germinal center formation. Regardless of these detailed mechanisms, it is clear infants can perform antibody selection as well as toddlers and adults, which provides some assurance of the effectiveness of vaccination in young children.

Another interesting finding is that infant antibody repertoires are capable of diversification, as seen through the clonal lineage analysis. We found that both infants and toddlers diversify their repertoire to the same degree upon an acute febrile malaria. Although adults have a larger range on the diversity of the lineage because they have accumulated more mutations, the degree of repertoire diversification in infants and toddlers is similar to what adults experience at an acute febrile malaria. This contrasts how elderly individuals preferentially increased the size of these lineages due to clonal expansion after influenza vaccination^6^. Together, these data provide evidence that infant antibody repertoires are similarly capable of responding to malaria as toddlers and adults.

Analyzing antibody lineages formed by sequences from both pre- and acute malaria samples, we were able to detect an increase in SHM upon malaria infection. While the high background SHMs at pre-malaria prevented us from detecting an increase of SHM upon malaria infection in the bulk repertoire except in IgM, our two-timepoint-shared lineage analysis showed both infants and toddlers increased SHMs upon acute malaria infection. This highlights the power of combining informatics analysis with improved technology – the amount of SHM increase in toddler was relatively small compared to infants, which would have been missed if molecular identifiers were not used. Similar problems may prevent a recent study from identifying an increase of SHM in healthy adults who received influenza vaccination^56^. It is also possible that the perturbation of a natural infection is much stronger than vaccination. The evidence we observed that infants have a similar propensity to diversify clonal lineages, perform antigen selection, and increase SHM load in acute malaria also provides a comprehensive assessment of the capacity of infants’ antibody repertoire to respond to a natural infection. These results also provided some assurance that infants are capable of responding to external stimuli and develop significantly diversified antibodies in response. These, along with the observation that administering anti-parasite drugs in the first 10 months of life correlated with higher malaria incidence in second year of life^57^, suggest that repeated vaccination might mimic the naturally acquired clinical malaria resistance that arises through repeated malaria exposures^26,57^.

Using progeny tracing in two-timepoint-shared lineages containing pre-malaria MemBs, we were able to detect total of 1799 lineages that contained these progenies. We analyzed conventional MemB cells because the atypical MemB frequency is low in this cohort of young children (**Supplementary Fig. 9**). This is possibly due to the relatively few malaria incidents in these young children, as opposed to adults and older children that we and others have analyzed where atypical MemB increases prevalence as a result of many years of chronic malaria exposure^58,59^. In depth analysis showed both continued SHM and isotype switching in MemBs in response to a natural infection rechallenge. Although we do not know the cell phenotype of these progeny sequences, it is reasonable to speculate that some of them could have developed into plasmablasts. Previous study has shown that plasmablasts have about 30 to 35 times more transcripts per cell compared to other peripheral B cell populations^60^. Even when we relaxed the threshold and counted all unique sequences that had more than 10 copies as plasmablasts, at most 2.1% of unique sequences in these pre-malaria MemB-derived progenies could be counted as plasmablasts (**Supplementary Fig. 16**). Applying the same threshold to bulk PBMC sequences from acute malaria timepoint resulted in a similar percentage of plasmablasts-derived sequences, which is consistent with about 2.2% of plasmablasts by flow cytometry analysis. All together, these data suggest that most of the progenies from pre-malaria MemBs remained as MemBs and continued to increase SHMs in these young children.

In summary, we presented an improved method of utilizing MIDs to significantly enhance the accuracy and coverage of the IR-seq. Using it, we systemically studied the antibody repertoire in malaria-exposed infants and toddlers and discovered several aspects of repertoire development, diversification, and capacity to respond to an infection that were not known before, which provides not only new parameters and approach in quantifying vaccine efficacy beyond traditional serological titer in the future but also venues for future studies of detailed molecular and cellular mechanisms of difference on antibody repertoire between infants and toddlers.

## METHODS

Methods are available in the Supplementary Information.

## ACKNOWLEDGEMENTS

The Authors would like to thank WE ARE BLOOD, blood center, for providing the blood samples for method validation; Dr. Scott Hunicke-Smith, Jessica Podnar, and Dr. Michael Wilson at the Genomic Sequencing and Analysis Facility at UT Austin for helping with the sequencing runs; Dr. Evan Cohen, for helping with the cell sorting; Dr. Purnamrita Sarkar at the Department of Statistics and Data Sciences at UT Austin for advices on statistical testing. Dr. Steven Kleinstein, Jason Vander Heiden, and Dr. Gur Yaari for sending us an updated version of the BASELINe software and advices on data analysis. This work was supported by NIH grants R00AG040149 (N.J.), S10OD020072 (N.J.), and R01GM106137 (P.R.) and by the Welch Foundation grant F1785 (N.J.). N.J. is a Cancer Prevention and Research Institute of Texas (CPRIT) Scholar and a Damon Runyon-Rachleff Innovator. B.S.W. is a recipient of the Thrust 2000 - George Sawyer Endowed Graduate Fellowship in Engineering. The cohort study in Mali was supported by the Division of Intramural Research, National Institute of Allergy and Infectious Diseases, National Institutes of Health.

## AUTHOR CONTRIBUTIONS

B.S.W. performed all malaria related experiment and data analysis; C.H. performed BASELINe and other sequencing data analysis; M.Q. developed the sequencing protocol using sorted NaiBs; D.W. helped with sequence analysis; S.M.H. helped with library construction; K.Y.M. helped with sequencing; J.X. helped with lineage visualization, E.W.L, P.D.C., and S.K.P. selected malaria subjects, provided samples and helped with experimental design; P.R. provided computation resources and helped with analysis; K.C. helped with lineage structure algorithm optimization and lineage visualization; N.J. conceived the idea, designed the study, directed data analysis, and wrote the paper with contributions from all co-authors.

